# Single-Neuron Encoding of Learnability in the Dorsal Anterior Cingulate Cortex

**DOI:** 10.1101/2025.09.29.679390

**Authors:** Yuhao Jin, Greg Jensen, Vincent Ferrera, Jacqueline Gottlieb

**Affiliations:** Department of Biological Sciences, Columbia University, New York, NY 10027; Department of Psychology, Reed College, Portland, OR 97202; Department of Neuroscience, Columbia University, New York, NY 10027; Kavli Institute for Brain Science, Columbia University, New York, NY 10027; Mortimer B. Zuckerman Mind Brain Behavior Institute, Columbia University, New York, NY 10027

**Keywords:** Learnability, Transitive inference, Anterior cingulate cortex, Monkey

## Abstract

In natural environments, associations that indicate true learnable regularities are intermixed with those that arise from random and ultimately unlearnable relationships between events. To efficiently allocate cognitive resources and avoid inferring spurious patterns, organisms must distinguish learnable from unlearnable associations, but the mechanisms underlying this ability are not understood. We recently showed that monkeys performing a transitive inference task, while discovering the true hidden order in learnable image sets, also behaved to varying degrees as if they inferred subjective order in objectively random (unlearnable) image sets. Here, we show that the ability to detect learnability is encoded by neurons in the dorsal anterior cingulate cortex (dACC, area 24c). dACC neurons responded strongly after a decision outcome as reported in previous studies and, additionally, signaled whether a trial was from a learnable vs unlearnable set before outcome delivery, and showed interactions whereby their selectivity for the outcome (reward vs lack of reward) was stronger for learnable versus unlearnable sets. Learnability and interaction responses were independent of sensory or reward cues (which were equated for learnable and unlearnable sets) but their strength correlated with the monkeys’ ability to avoid inferring false order in unlearnable sets. The findings suggest that the dACC is part of a network that monitors learnability and enables animals to appropriately focus learning on true patterns while avoiding false inferences about spurious and random associations.

## Introduction

Cognitive neuroscience has made significant progress in understanding the theoretical and neural basis of associative and rule-based learning in environments in which participants are exposed to predictable rules, and are rewarded for learning responses that conform to these rules. Such environments, however, differ greatly from natural settings, in which animals face not only consistent associations that can be fruitfully learned but also many coincidental associations that are random and unlearnable. To learn efficiently in such settings individuals need not only learn but *decide when to learn* (i.e., distinguish true from random associations and direct learning accordingly). This ability is crucial both to avoid laboring in vain on ultimately unlearnable tasks, and to avoid erroneously inferring fictitious structures in objectively random events. Despite its importance, however, the behavioral and neural mechanisms of learnability detection are not well understood.

Using learnability for controlling behavior poses a considerable computational challenge because it requires animals to estimate whether a true structure exists before fully learning the structure. In experiments that provide explicit learnability cues, participants readily react to these cues (e.g., humans modulate evidence accumulation and learning rates depending on whether they are instructed that rewards are random or that rewards are action-contingent, learnable and controllable)^1,2^, and mice slow their choices to boost learning earlier in a session, but only if they view discriminable (learnable) and not blurred and non-discriminable visual cues^3^. However, studies using more ecological settings in which learnable and unlearnable stimuli are randomly interleaved and uncued produced mixed results. On one hand, infants attend more to unlabeled learnable vs random linguistic grammars^4^ and pigeons prefer otherwise identical sequences that differ only in the presence or absence of consistent relationships between stimuli and rewards^5^. On the other hand, humans who could freely allocate study time tended to focus on unlearnable games at the expense of mastering learnable games^6^, and humans often suffer from illusions of control: the erroneous belief that one can predict stochastic events^7–9^. Thus, humans and animals show variable capacity to autonomously estimate learnability in natural settings, which may depend on the specific individual, environment and task.

A crucial question concerns the neural basis of learnability detection. A recent computational model, building on the proposed roles of the anterior cingulate cortex (ACC) in in monitoring and control^14–18^, proposes that the ACC acts as a “meta reinforcement learning controller” that recruits cognitive processes (attention, memory and/or learning rates) based on the context and demands of a task^11–13^. Consistent with this view, studies using fMRI and ERP have shown that the human executive network is sensitive to controllability when it is explicitly cued^1,2,8–10^. It remains unclear, however, whether or how this network encodes autonomous estimates of learnability independently of such cues.

Here we examined this question by recording dACC neuronal activity during a transitive inference (TI) task in which monkeys learned to recognize abstract structures in ordered versus random pictorial sets^19,20^. In each session, monkeys made choices between image pairs that could be drawn from one of two sets: a learnable (L) set in which choices were rewarded according to a consistent hierarchical order (i.e., A was better than B, B was better than C, etc.), or an unlearnable (U) set in which choices were rewarded randomly. Importantly, L and U trials were fully unsignaled, being randomly interleaved and equated in terms of appearance and reward rates, and differed only in the presence or absence of a latent hierarchical structure. We recently showed that, while monkeys reliably discovered the true hierarchy of the L sets, they often chose as if they inferred a subjective order in the U sets, and this behavior reflected the use of an internal model of real or fictitious ordering that contravened simple reward-based associative learning^21^. Here, we show that individual ACC neurons encode learnability independently of rewards and correlate with variability in the monkeys’ ability to distinguish L and U sets. The findings illuminate the neural mechanisms by which animals autonomously estimate learnability and avoid inferring fictitious structures in random events.

## Results

### Monkeys showed variable ability to distinguish between learnable and unlearnable sets

Adult male rhesus monkeys (*N*=2) performed a task in which they chose one of two pictorial stimuli and were rewarded either randomly or according to a learnable rule. Each trial began with 500 msec of central fixation, after which the monkeys were shown two images at symmetric locations in the right and left visual field and, after a 400 ms delay, made a saccade to the stimulus of their choice (**Fig. 1A, B**). During each behavioral session, monkeys were tested with 20 novel image pairs organized in two sets. Images in the “learnable” (L) set had an objective (hidden) ranking (**Fig. 1C**) and monkeys were rewarded if they chose the item with the superior rank in each pair (e.g., given image pair B, D, reward was delivered for B but not D; **Fig. 1A**). In contrast, images in the unlearnable (U) set had no objective ordering (**Fig. 1D**), and monkeys were rewarded probabilistically and independently of their choices (**Fig. 1B**). Each session introduced new L and U pictorial sets, and monkeys interacted with them over a minimum of 784 trials, including an initial training phase of 384 trials presenting only 4 adjacent-rank pairs from the L set and 4 randomly chosen pairs from the U set, and a subsequent testing phase randomly interleaving all 20 possible pairs from both sets (**Fig. 1E, F**). Monkeys G and F completed, respectively, 40 and 41 sessions with concurrent recording of dACC cells.

**Fig. 1.**
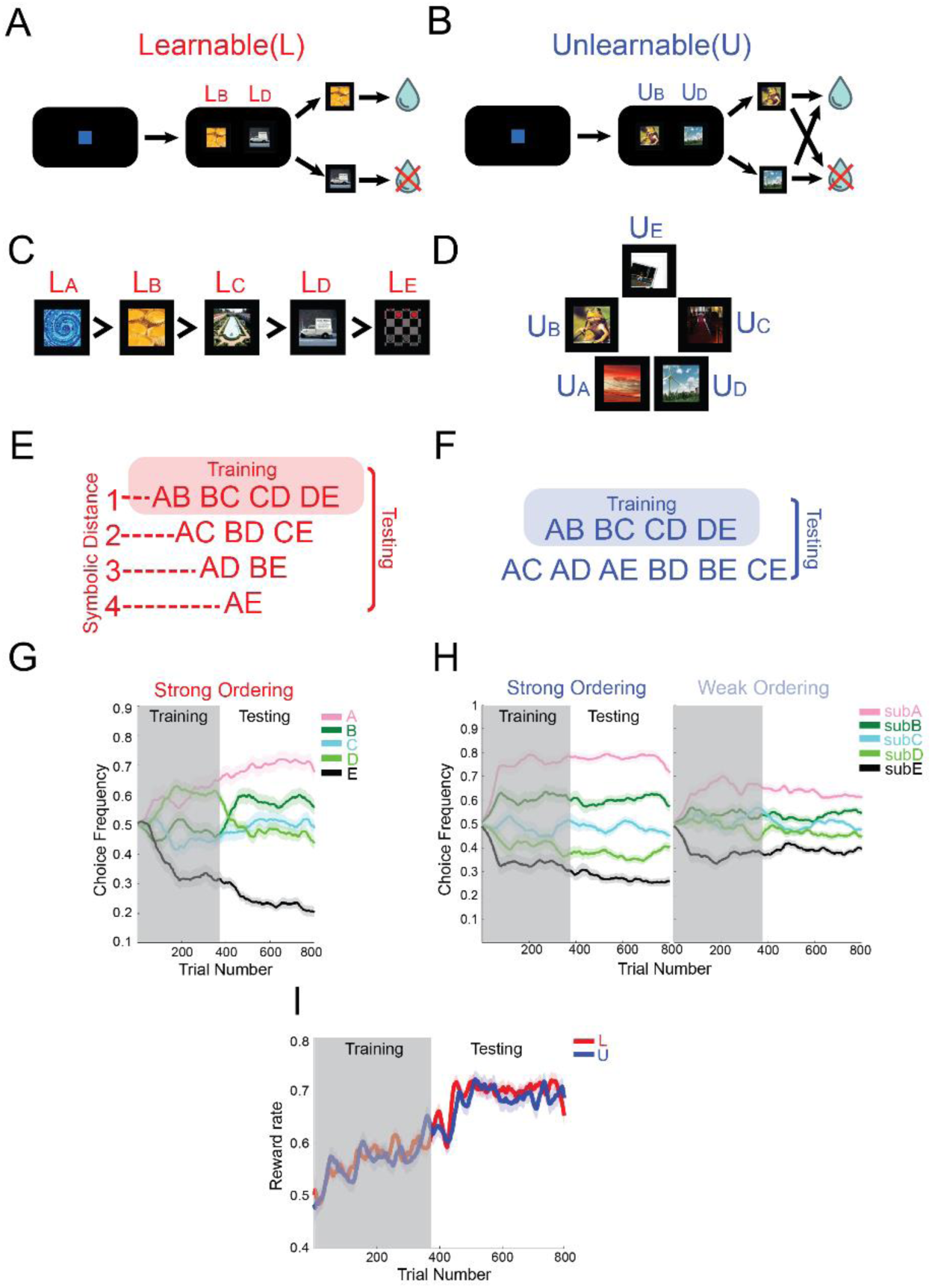
Task A-F: task paradigm. **A-D**: Task consisted of two pictorial lists with five items each where one list faithfully observed a predictable order (Learnable L list, **C**) whereas the other was completely unordered (Unlearnable U list **D**). Outcome was totally hinged or independent on the relative ordering for the L and U list separately. **E-F**: The task was divided into training phase (only the adjacent pairs in L and four random pairs in U) followed by testing phase with all the pairs presented. **G, H:** the evolution of preference levels over all stimuli over trials for the L set (**Fig G left, according to the objective rank**), strong ordering U set (**Fig H left**) and weak ordering U set (**Fig H right, according to the subjective rank**), which was calculated as the sliding window averages of chosen frequencies (n= 32 and 40 for the training and testing, stepped by 1 trial). **I**: The evolution of reward rates across trials. Traces show the reward rates for L (red) and U (blue) stimuli as a function of trials (sliding window of 10 trials stepped by 1 trial, N = 40 in monkey F and N = 41 in monkey G). Reward rates on the L set increased across trials, indicating the monkeys’ learning the correct ordering, and were dynamically yoked with those on the U set as explained in the text. Error bars in **G-I** denote standard error (SE).

### Monkeys showed subjective ordering on the U set whose strength differed by session

Consistent with our previous report on this task^21^, the monkeys reliably learned the correct ordering for the L sets during the training phase and quickly generalized it to the new pairs at the start of the testing phase (**Fig. 1G**). Importantly, the monkeys also developed robust ordering preferences on the objectively random U set, consistently choosing as if some stimuli had higher ranks relative to others, although the strength of this preference differed by session (**Fig. 1H**). These preferences could not be ascribed to visual cues, as images from the L and U sets were visually equivalent and randomly interleaved. Importantly, we dynamically adjusted the reward probability on the U set to be equal to that on the previous 10 trials on the L set, and verified that the reward rates were identical on the two sets, both on average and in their trial by trial dynamics (**Fig. 1I**, Comparison of averaged reward rates between L and U, Wilcoxon test, Pooled: *p* = 0.68; Monkey F: *p* = 0.72; Monkey G: *p* = 0.74). Thus, reward rates provided no explicit cues to the ordering that was in effect in a trial.

As in our previous study^21^, we quantified the monkeys’ ordering preferences (OP), by fitting choice data during the testing phase with a hierarchical Bayesian model that assumed a noisy internal representation of stimulus rank (**Fig. 2A**, top; ***Methods***). Using this fitting procedure, we derived a *z*-score indicating the monkeys’ preference for each particular image, such that more dissimilar/similar *z*-scores indicated stronger/weaker OP for a particular set (**Fig. 2A**, bottom; ***Methods***). We then plotted the *z*-scores as a function of stimulus rank (the objective rank for the L set and subjective rank for the U set) and used the slope of the linear fit as a measure of OP (respectively, OP_L_ and OP_U_; **Fig. 2B**), to determine if the empirically measured OP exceeded the baseline expected from random responding (***Methods***).

**Fig. 2.**
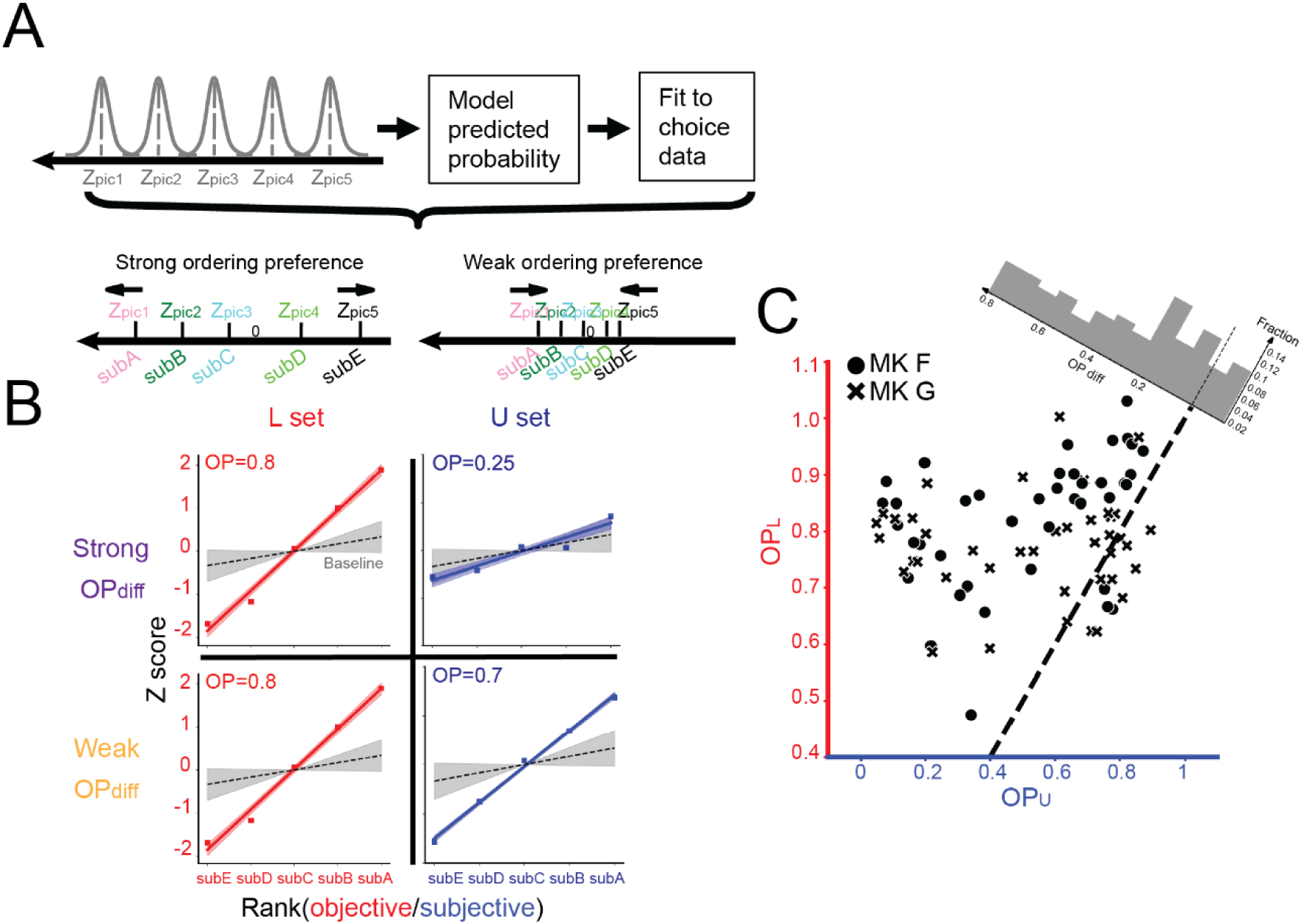
Analysis of ordering preference. **A. Pipeline for subjective ordering analysis.** The chosen frequencies for each stimulus were fit by a mechanistic model assuming that stimuli had an ordered internal representation, which produced a set of *z*-scores indicating weaker (**left**) or stronger (**right**) subjective ordering. **B**. **Example sessions with strong and weak OP_Diff_.** The top and bottom rows show two representative sessions with, respectively, strong versus weak behavioral differentiation between L and U sets. In each session, the colored numbers show the OP for the L and U sets, and the colored traces show the corresponding linear fits of *z*-scores as a function of rank. Shading shows 95% CI of the fitted slopes. Gray traces indicate baseline slopes inferred by simulating an agent that emits random responses (average and 95% CI of fitted slopes over 81 simulations). **C. Comparison of OP_L_ and OP_U_.** Each point is one session. The dashed trace is the equality diagonal. The gray distribution shows OP_Diff_. Within it, the dashed trace indicates equal ordering for L and U sets (OP_Diff_ = 0), and positive values (plotted to the left) indicate greater behavioral differentiation.

OP_L_ values had an overall mean and standard error (SE) of 0.79 +/- 0.012, which was significantly greater than the baseline OP in each monkey (**Fig 2B**; monkey F: 0.82 +/- 0.018, *p* < 0.0001 vs baseline of 0.18 +/- 0.016; N = 41, paired t-test; monkey G, 0.77 +/- 0.014, *p* < 0.0001 vs baseline of 0.172 +/- 0.014, N = 40, paired t-test). OP_L_ values were significantly above baseline in each individual session (95% CI monkey F: [0.79,0.86]; monkey G: [0.75,0.8]), confirming that both monkeys reliably inferred the true order of all the L sets.

Remarkably, OP was also greater than expected by chance for U sets, indicating that monkeys had a significant tendency to respond as if these sets had a hidden order (OP_U_, monkey F: 0.51 +/- 0.04, p < 0.0001 vs baseline of 0.18 +/- 0.016, N = 41, paired t-test; monkey G, 0.53 +/- 0.04, p < 0.0001 vs baseline of 0.172 +/- 0.014). However, OP_U_ was more variable relative to OP_L_ (standard deviations of 0.27 vs 0.10; Fligner-Killeen Test, N = 81, *p* < 0.0001) suggesting that subjective ordering preferences for the U sets varied by session. Because we were interested in the monkeys’ ability to distinguish between the two sets, we computed the differential OP (OP_Diff_) by taking the difference between OP_L_ and OP_U_ for each session. As shown by the histogram in **Fig. 2C**, OP_Diff_ formed a broad distribution ranging from values close to 1.0 (indicating sessions in which the monkeys showed clear ordering for the L but not U sets) to near-zero for sessions in which monkeys treated both sets as being equally ordered.

### dACC Neurons Encode Learnability Independently of Reward

To examine the neural mechanisms underlying the detection of learnability, we recorded neural activity in the dorsal anterior cingulate sulcus (dACC), focusing on the dorsal portion of the anterior third of the sulcus (Brodmann area 24c) that has been reported to respond to uncertainty and conflict^22,23^ (**Fig 3A**; ***Methods***). We report the activity of 1072 neurons (436 from monkey F) that were recorded using multi-contact probes and classified as single-units based on offline sorting, but were not otherwise pre-selected for specific task responses (***Methods***). The neurons had sustained elevated responses between the onset of the choice stimuli until after the monkeys’ saccades, and additional phasic responses to the delivery of the outcome (reward or no reward accompanied by, respectively, high and low pitch tones; **Fig 3B**).

**Fig 3.**
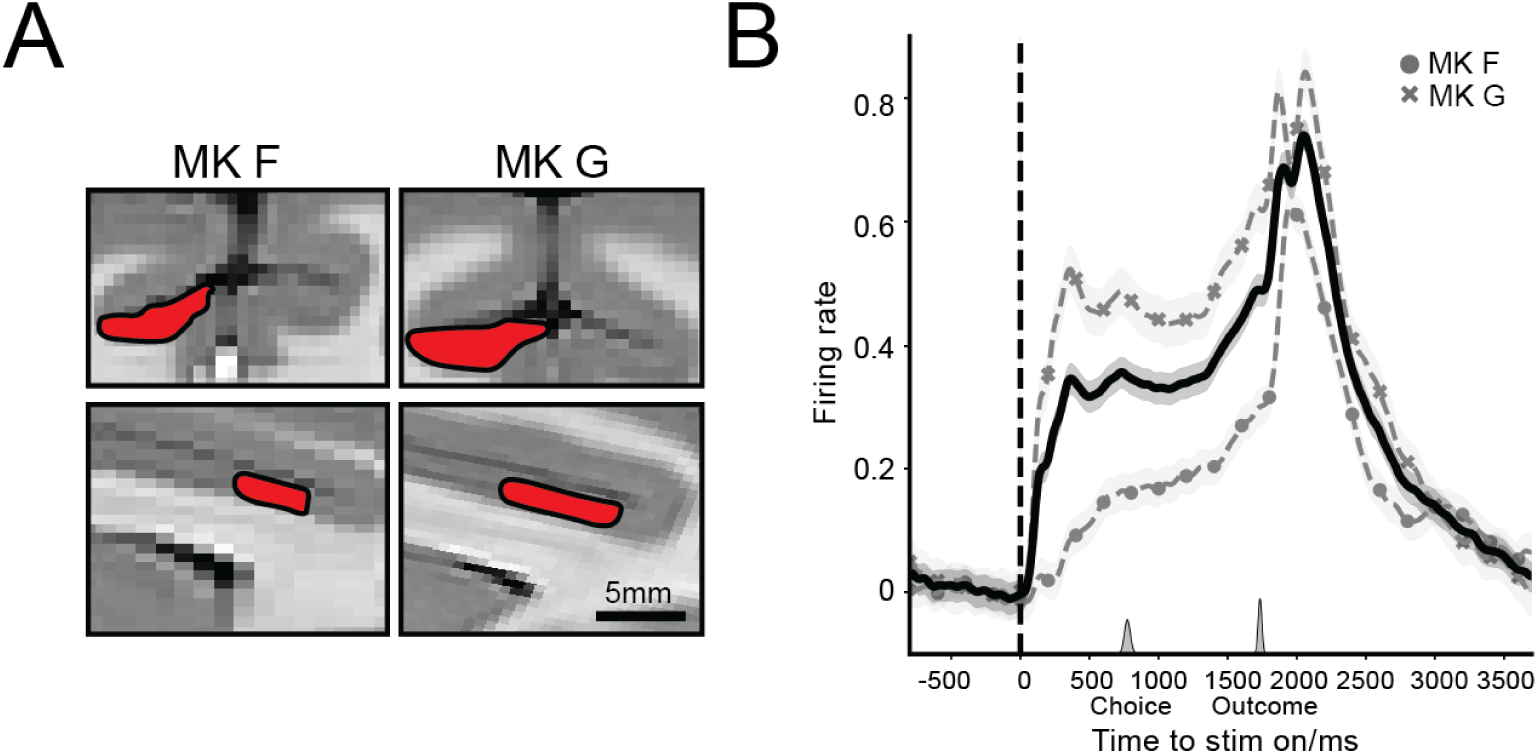
Summary of recording sites and neural response. **A. fMRI images showing the approximate extent of the recording areas for each monkey.** The top images show the coronal plane 26 mm anterior to anterior commissure in each monkey; the bottom images show the sagittal plane, 3.5 mm to the left of midline. **B. Average neural response aligned on stimulus onset.** Traces show mean and SE of the peri-stimulus time histograms across all neurons for each subject (Monkey F: circle, N = 436 neurons; Monkey G: cross, N = 636 neurons) as well as combined (solid. N = 1072). The small gray histograms on the x-axis show the distributions of saccade onsets and outcomes delivery across all trials.

To determine whether and how the cells encoded task variables, we fit firing rates using a GLM model that included trial-by-trial regressors for learnability (L vs U), outcome (reward/no-reward) and their interaction, alongside a nuisance regressor for saccade direction to rule out directional selectivity confounds, and evaluated the model for each cell in 3 task epochs (800 ms pre-choice, post-choice and post-outcome; ***Methods***).

Over one third of the cells (36.8% (395/1072); monkey F, 47% (205/436); monkey G, 30% (190/636)) significantly encoded learnability in at least one task epoch (Wald z test, *p* < 0.05 after Benjamini-Hochberg multiple comparison correction). Of this subset, slightly over half (monkey F: 53% (108/205); monkey G, 58%, (111/190)), had positive coefficients indicating stronger responses for L vs U sets, while the remainder had significantly stronger responses for U vs L sets, as illustrated in **Fig. 4A**, left (top vs bottom). To estimate the time-course and strength of the learnability effects, we fit the GLM model (***Methods***) to time-resolved firing rates (average firing rate in 300 ms moving windows with a 50 ms step) and computed the coefficient of partial determination (CPD), which measures the fraction of each neuron’s firing rate variability attributable to each factor (***Methods***). Learnability accounted for a significantly larger fraction of variance than expected by chance (after shuffling trial labels; ***Methods***) throughout the pre-choice, post-choice and post-outcome epochs. This was the case in both the subset of responsive cells (**Fig. 4B**, left, thin trace) and across the entire population of cells (**Fig. 4B**, left, thick trace).

**Fig 4.**
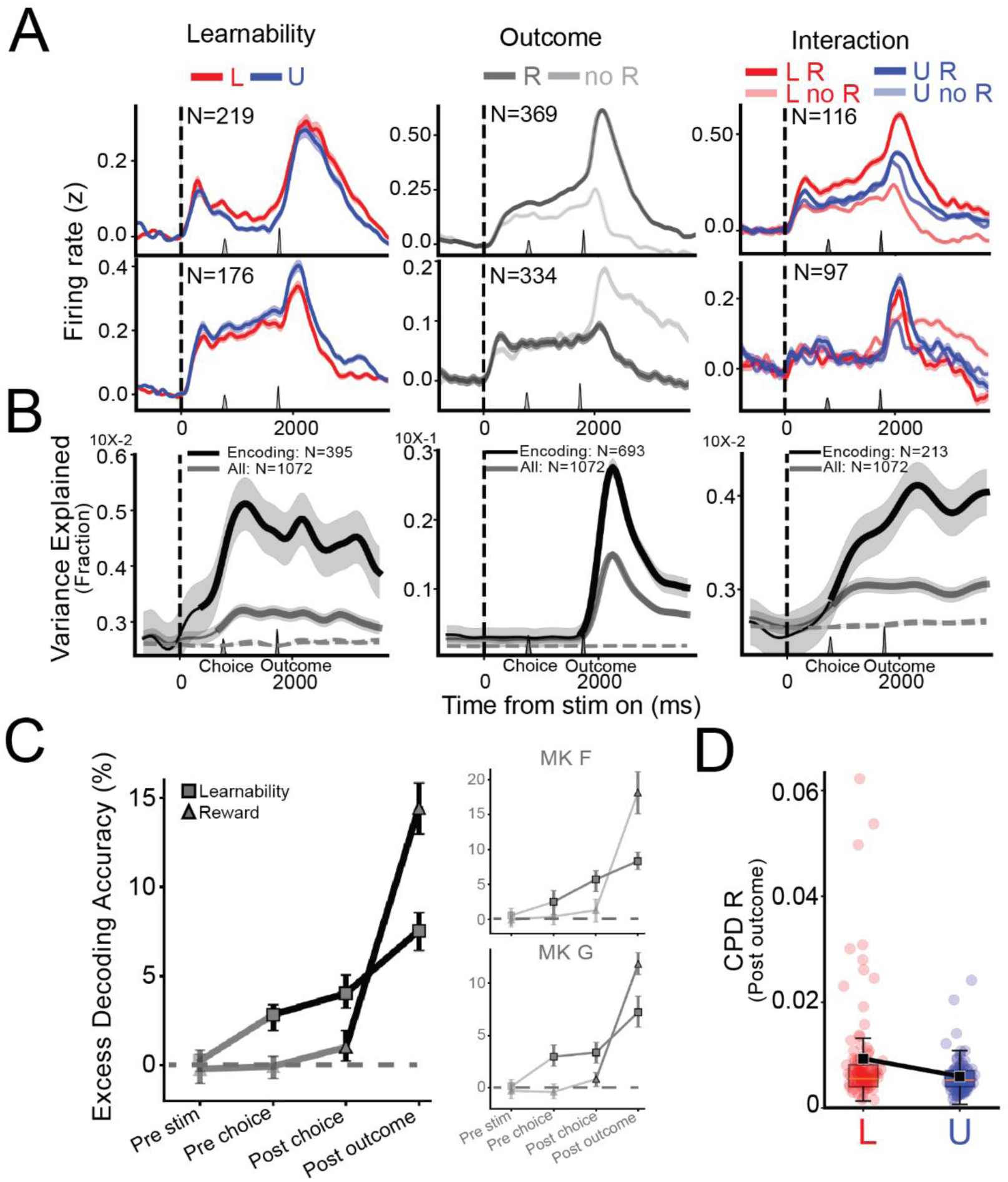
dACC neurons encoded learnability, outcome and their interactions. **A**. **PSTHs for the subsets of neurons encoding Learnability, Outcome and their Interactions.** Neurons are separated by the signs of the GLM coefficients as explained in the text. Each trace shows the average and SE of the PSTH over the indicated number of cells (constructed after z-scoring relative to the averaged firing rates over 500ms before stimulus onset). Other conventions as in Fig. 3B. **B**. **CPD for Learnability, Outcome and Interaction.** Each trace shows the mean and SE of the CPD computed in a sliding window (size = 300ms, stride = 50ms) and averaged over the cells encoding the respective variable (black trace) or the entire population of cells (dark gray trace). Thick points in each trace show windows in which the CPD differed from the shuffled control. Dashed traces show the shuffled control (mean and SE over 50 iterations). **C. Cross-validated decoding accuracy of Learnability and Outcome**. The traces show the average of the excess decoding accuracy, defined as the accuracy in decoding each variable based on the firing rates of all cells minus the average accuracy after shuffling trial labels. Error bars show 95% CI, and darker symbols show significant decoding (CI’s do not overlap 0). The large panel on the left shows the pooled data and the smaller panels on the right show individual monkeys. **D. CPD for outcome encoding is larger in L vs U sets.** Each point is the averaged CPD for the outcome (reward vs no reward) across neurons for each session and set. A box-and-whisker plot is overlayed on the points; large black symbols show mean and SE.

Independently of their learnability modulation, nearly 2/3 of the cells encoded the outcome, distinguishing between a reward and lack of reward (Wald *z* test, *p* < 0.05, Benjamini-Hochberg correction; 66% (^703^/_1072_) overall; monkey F, 65% (^282^/_436_), monkey G, 66%, (^421^/_636_)). Among the reward-encoding cells, similar fractions showed positive and negative coefficients, responding significantly more strongly after a reward vs no-reward outcome (overall: 53% (^369^/_703_); monkey F 52% (^146^/_282_), monkey G 53%, (^223^/_421_)), with the remainder showing the opposite modulation as depicted in the top and bottom rows in **Fig. 4A** (middle column). Interestingly, some positive reward-coding was present before reward delivery (**Fig. 4A**, top row middle column) reflecting the fact that, on L sets, the reward outcome was strongly associated with the rank of the chosen stimulus and could thus be predicted before the choice (***Methods***). However, time-resolved CPD analysis showed that the fraction of variance explained by the outcome was significant only after outcome delivery, in both the reward-responsive cells and the whole population (**Fig. 4B**, middle column).

Together, the results show that dACC neurons had robust responses to learnability that were distinct from their previously-documented outcome responses. To verify the robustness of this finding and better understand the relative time-course of the two signals, we used the alternative method of logistic decoding to classify each variable based on the firing rates of all cells. Cross-validated decoding accuracy (in sessions with more than 10 simultaneously recorded cells) significantly exceeded the expected null-level performance for both learnability and outcome, showing that both variables could be reliably decoded from the overall dataset and each individual monkey (**Fig. 4C**). In the pre-choice, post-choice and post-outcome epochs, learnability was decoded with average accuracy of, respectively, 54.0% [53%, 54.6%], 55.1% [54.3%, 56%], and 58.3% [57.2%, 59.3%],significantly exceeding the levels expected by chance in each case (respectively, xx% [51%, 51.2%], xx% [51%, 51.2%], and xx% [50.7%, 50.9%]). Interestingly, learnability decoding persisted and even increased in post-choice and reward vs the pre-choice epochs, suggesting that the neurons conveyed sustained signals of learnability rather than merely distinguished the L vs U image sets. Moreover, while learnability was decoded before the monkey’s decision, reward decoding became marginally significant only after the decision and was strong only after outcome delivery (**Fig. 4C**).

Given that the neurons encoded both learnability and the outcome, we examined if these responses interacted – i.e., if learnability modulated the neurons’ ability to distinguish between a reward and no-reward outcome. Consistent with this hypothesis, the GLM analysis showed that 20% the cells had significant interaction coefficients in at least one task epoch (Wald test, *p* < 0.05, Benjamini-Hochberg corrected; overall: 20%, (^213^/_1072_); monkey F: 19%, (^85^/_436_); monkey G: 20%, (^128^/_626_)). Interaction coefficients could be positive or negative based on whether the cells responded more to a reward or lack of reward, as noted above (**Fig. 4A**, right). Time-resolved CPD analysis showed that interaction effects accounted for a significant fraction of firing rate variance during the post-choice and post-outcome epochs, both in the subset of sensitive cells and in the full population (**Fig. 4B**, right panel).

Interestingly, whether the coefficients were positive or negative, the interactions reflected a stronger neural differentiation between reward and no-reward outcomes on L than U sets. Thus, neurons with positive interactions showed a stronger preference for reward over lack of reward in L vs U sets (**Fig. 4A**, top right), and neurons with negative interactions showed a stronger preference for no-reward vs reward outcomes in L vs U sets (**Fig. 4A**, bottom right). This was confirmed by a follow-up analysis in which we computed the CPD for encoding reward/no-reward during the post-outcome epoch for each stimulus set, and found that CPD was significantly larger in L vs U sets (Fig. **4D**, ***Methods***, pooled data *p* = 0.04, monkey F, *p* < 0.0001; monkey G, *p* = 0.05, Wilcoxon rank-sum test).

In sum, dACC neurons encoded learnability independently of rewards and showed stronger reward-related responses in L vs U sets.

### Neural Learnability and Interaction Effects Correlate with Behavior

To determine how the neural responses were related to behavior, we examined the correlations between the strength of neuronal encoding the monkeys’ ability to show differential ordering of L vs U lists (denoted by OP_diff_). We reasoned that, if the neurons contributed to behavior, on sessions in which the monkey was better able to differentiate between the L and U lists, the neurons would also show stronger encoding of Learnability, Outcome and/or Interactions.

We found that OP_diff_ showed significant positive correlations with the neural encoding of Learnability and Interaction but not Outcome. **Fig. 5A** illustrates the results from a session-level analysis in which we correlated the OP_diff_ for each session with the average CPD across the cells in that session. Spearman rho values were significantly positive for Learnability (0.29, *p* = 0.01), and Learnability x Outcome interaction (0.41, *p* = 0.0002) but not for Outcome CPD by itself -0.02, *p* = 0.88). The results were robust in each monkey individually (monkey F: respective rho = 0.32 (*p* = 0.05), 0.35 (*p* = 0.02) and -0.11 (*p* = 0.48); monkey G 0.39 (*p* = 0.013), 0.58 (*p <* 0.0001) and 0.06, *p* = 0.69). Results were also robust in an alternative analysis that correlated session-level OP_diff_ with the CPD for individual cells in a session (full dataset, rho = 0.2 (*p* < 0.0001), 0.15 (*p* < 0.0001) and 0.11 (*p* = 0.12) and for, respectively, Learnability, Interaction and Outcome, replicated in each monkey).

**Fig 5.**
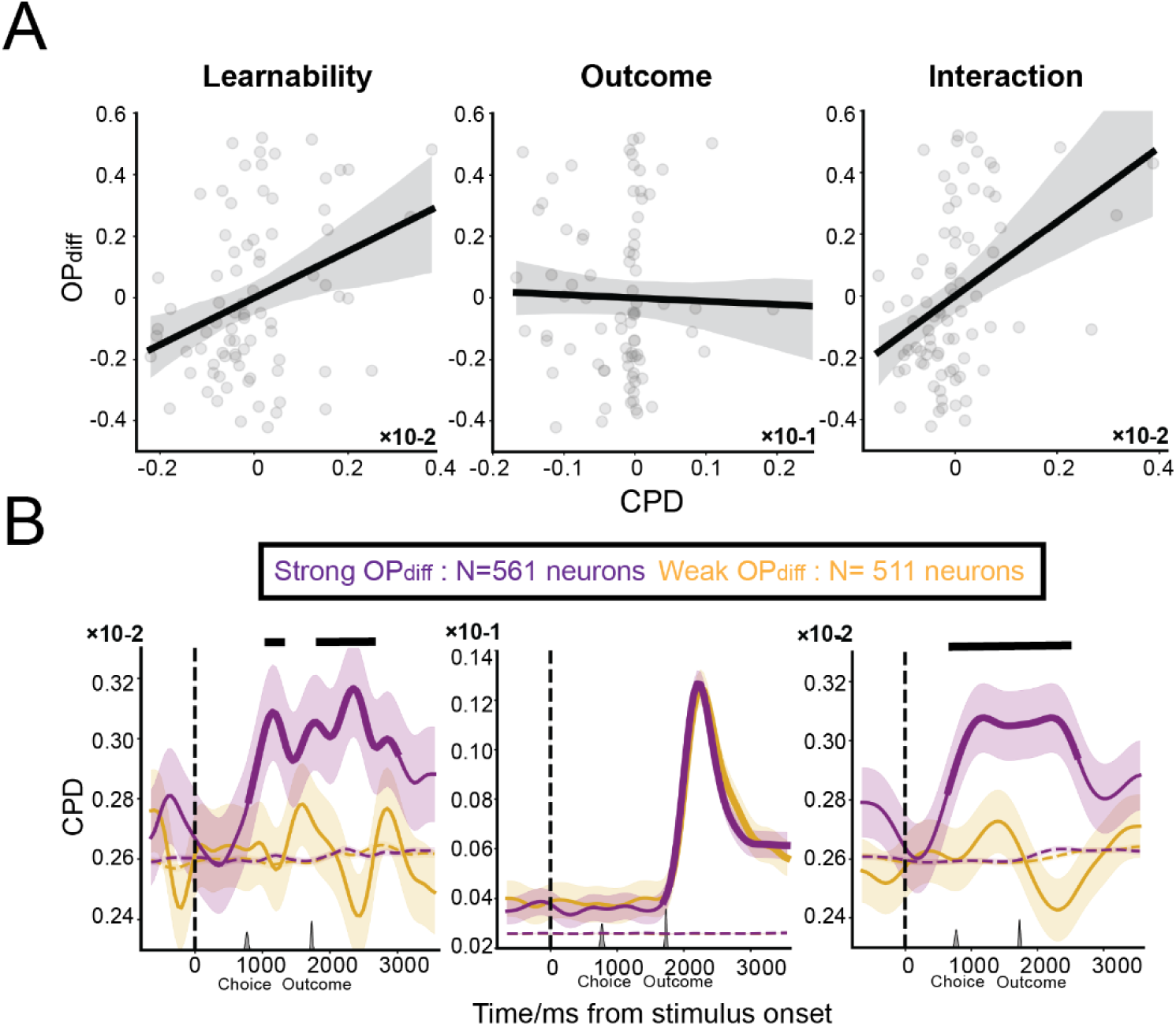
dACC activity correlates with behavior. **A**: **OP_diff_ correlates with CPD for Learnability and Interaction, not Outcome.** Each point shows the OP_diff_ for one session (abscissa) and the CPD for the respective variable, measured in the post-outcome epoch and averaged across all the neurons recorded in that session (ordinate). The lines and shading show the best fitted regression line and its 95% CI over *N* = 81 sessions. **B**. **CPD for Learnability and Interaction, not Outcome, differed between sessions with above-median vs below-median OP_diff_** (respectively, purple and yellow). The traces show mean and SE of the time-resolved CPD across the neurons in each group (*N* indicated in the legend above). Dashed horizontal traces show the averaged CPD with each corresponding labels shuffled 50 times. Thicker points show time bins with CPD significantly different from its own shuffled control and horizontal bars at the top show whether CPD differed between the two groups by independent *t*-test. All other conventions as in Fig. 3B.

To illustrate the time-course of the behavioral differentiation, we plotted the time-resolved CPD on sessions with above- and below-median OP_diff_ (**Fig. 2C**; performing the median split within monkey to rule out subject-specific confounds). As shown in **Fig 5B** (middle panel) CPD for the Outcome was high and indistinguishable in the two behavior groups. In contrast, CPD for Learnability and Interactions was near-chance in sessions with below-median OP_diff_, was significantly stronger, and was significantly above-chance in sessions with below-median OP_diff_ throughout the course of the trial **Fig 5B** (right and left panels).

Thus, the dACC responses to learnability, and the differential encoding of the outcome for L vs U sets, were significantly correlated with the monkeys’ ability to distinguish between the two sets.

## Discussion

Natural environments contain a practically infinite number of potential associations between disparate features and events. However, only a tiny minority of these associations signal consistent, learnable structures, while the vast majority are spurious and unlearnable. Distinguishing learnable from unlearnable contexts is therefore crucial, both to avoid laboring in vain and to prevent erroneous inferences about the presence of structures in objectively random events. Using learnability to control learning, however, is a significant challenge, because true and random associations are not labeled as such, and animals must infer hidden differences between these sets of associations before fully learning the structure.

Here, we show that monkeys have some – albeit variable and imperfect – ability to distinguish structured vs unstructured pictorial sets, and crucially, this ability is correlated with the ability of dACC cells to distinguish these contexts. We show that individual dACC neurons and populations of cells encoded learnability independently of reward outcomes, and these responses varied in strength in a manner that was correlated with the monkeys’ ability to differentially learn in L versus U sets. Regression analysis, combined with the precisely-matched reward rates on L and U sets, showed that the learnability responses were clearly distinct from the trial’s rewards, and thus could not be explained by quantities such as reward prediction errors/surprise^24^, reward uncertainty^25^ or reward expectation^26^ that had been shown to be conveyed by dACC cells.

The neural encoding of learnability persisted after the monkey’s decisions and interacted with outcome-related responses, such that the neuronal differentiation between a reward and a lack of reward was stronger in L vs U sets. This result is consistent with a substantial body of evidence from fMRI and ERP studies in humans showing reward-related responses are dampened in uncontrollable vs controllable tasks^1,34,35^. In our task, the dampened response to reward on one trial may have reduced the extent to which monkeys adjusted their choices on the following trial. Thus, in sessions in which this differential dampening was stronger (i.e., with stronger interactions between learnability and outcome-related responses) animals may have been better able to learn the true structure based on reward feedback on L sets but ignore the feedback on U sets, explaining the correlations with OP_diff_ we found in the data. This possibility, as well as how it such an incremental reward-based mechanism may interact with internal representations, can be fruitfully explored in future experiments using detailed computational modeling of behavior in humans and monkeys.

Our results are consistent with a role of the dACC in executive control and monitoring^14–16,27,28^, and with a neurocomputational model of meta-level control of attention and learning based on the task context and goals^12,13^. However, our findings suggest that executive regulation relies on more than external rewards or sensory cues, as has been assumed in previous work. In our task, the rates of reward on L and U sets were dynamically and precisely equated. Thus, L and U sets could not be distinguished in terms of previously proposed reward metrics, including the rates of reward, the uncertainty (volatility) of these rates^12,13^, or learning progress defined as the derivative of the rates over time^6^. Similarly, learnability responses could not be explained by visual cues. While the animals may have detected the presence of two pictorial sets (i.e., the fact that images from L and U sets did not co-occur in a trial), this cannot explain the dACC learnability responses, which persisted and grew after the choices (when the images disappeared) and, crucially, correlated with behavioral preferences. Our findings thus suggest that the dACC detects learnability based on mechanisms that are independent of both external rewards and sensory cues.

What might be the nature of these mechanisms? One possibility is that the learnability responses in the dACC reflected, at least in part, the monitoring of internal representations of a hierarchical structure. This is consistent with the evidence that such representations, whether they reflected a veridical structure on an L set or a fictitious structure on a U set, were necessary to explain behavioral choices^29,30^. An alternative possibility is that the monkeys inferred learnability based on prior beliefs, simply assuming that U sets have a learnable order based on their prior experience with various learnable tasks. A final, not mutually exclusive possibility, is that animals monitor the predictability between simulated operations and corresponding outcomes (e.g., the mutual information between their choices and rewards). This quantity is a measure of “empowerment” in agent-centric learning and AI^31–33^, and is emerging as a strong candidate for a type of intrinsic motivation that guides learning independent of local reward. Thus, a critical question for future research concern the types of intrinsic motivations – be they based on prior beliefs, internal representations or empowerment – that contribute to the regulation of learning and other cognitive functions.

In sum, our results show that dACC neurons detect learnability independently of sensory cues and rewards, and may underlie the intrinsically-motivated control of learning in complex environments containing unknown mixtures of structured and random associations.

## Materials and Methods

### Subjects

We trained two adult male rhesus macaques (*Macaca mulatta*, F and G, weighed 8.1 and 10 kg during the data collection) for our task. Besides basic training on eye fixation and visual guided saccades, subject F had prior training on the transitive inference task but neither was pretrained on our dual list paradigm. Subjects collected liquid rewards from the task. Our study was approved by the Institutional Animal Care and Use Committees (IACUCs) at Columbia University. Research was also conducted according to the guidelines from the Guide for the Care and Use of Laboratory Animals of the National Institutes of Health (NIH).

### Behavior control

We implanted each subject with a metal bar over the acrylic headpost situated around the posterior midline, by which they were head fixed in the primate chair while interacting with the task by making the eye saccades. In order to infer the monkey’s eye position on the screen, we used an infrared camera for capturing video of one eye (Flea 3 FL3-U3, Point Grey, Wilsonville, OR, 600Hz frame rates), after which software (fly capture, FLIR, Richmond, British Columbia, Canada) automatically detected the center and outline of eyes, and output the *x* and *y* coordinates. Before the experiments, eye position was calibrated by requiring subjects to fixate on the small white square in the center and perimeters of the display for 500ms, during which we aligned monkeys’ eyes with the square locations. Stimuli were displayed by a CRS VSG 2/3F video frame buffer on an CRT high resolution monitor (sample frequency 60Hz, 1280×1024 pixels). We sized each stimulus at 140×140 pixels, located them symmetrically around the center (eccentricity of 200 pixels from the picture center) and set the viewing distance at 60cm. After the subject made the saccade, eye coordinates were sent to the behavior computer that proceeded to the next phase of the trial. To deliver the reward, the computer sent out two pulse signals to a solenoid valve resulting in two drops of water (0.1cc each) being pumped through the sipper tubes installed on the primate chair. We mounted another camera 45 degrees left to the chair and recorded the monkey’s face during the whole session by which we would be able to extract the time and frequency of licking each trial for future use.

### Task design

In each session, 10 pictorial stimuli were selected from a database of over 2500 images and made sure they were not presented before. Those 10 images were randomly assigned to one of the two lists with 5 stimuli each. Of the 2 lists, one was learnable (L) where each stimulus was arbitrarily assigned with its unique rank and subjects were required to deduct the veridical order by trial and error. The other set was unlearnable (U), meaning pictures within were unordered by which subjects couldn’t obtain any predictable relationships among stimuli under such context.

Each trial started with a small solid white square appearing at the center of the screen, after which the monkey was required to move his eyes towards it within 1500ms. After that they had to fixate on the square for another 500ms to proceed with the trial. Not fixating in time or breaking from fixation aborted the trial immediately. On average, monkeys broke about 25% of trials (Finish rate, Monkey F: 0.8 ± 0.03; Monkey G: 0.71± 0.04) each session suggesting good motivations. After the fixation period, 2 pictures were randomly drawn from one list and appeared equidistant from the screen center in opposite directions. Within three seconds, monkeys needed to respond by making a visual saccade to the left or right picture and hold their eyes in the acceptance window of the picture (150 by 150 pixels square) for another 500ms to confirm the choice.

Feedback depended on the trial learnability. If the pairs were from the L set, the subject received two drops of water and simultaneous high frequency tone (880 Hz) if the choice followed the correct order or two seconds of screen being completely dark as the punishment as well as low frequency tone otherwise (440 Hz). On the other hand, if the trial was unlearnable, feedback was independent of subject’s choice so that reward probability for choosing either stimulus was equal to the mean reward rate over the past ten learnable trials relative to the current trial.

Sessions were composed of training blocks and testing blocks. Each training block contained 48 trials, including four adjacent pairs (AB, BC, CD, DE) from two stimulus lists and each pair were presented 6 times with spatial counterbalance applied. Testing blocks followed a similar structure, presenting each of all 10 possible stimulus pairs for the two lists, each presented twice to counterbalance their positions on screen, resulting in 40 trials every block. L and U trials were interleaved by randomly shuffling the presenting sequence to keep the animal from predicting the trial learnability and certain image pair. Both monkeys needed to complete eight training blocks before proceeding to the testing block. They were allowed to finish as many testing blocks as they could until satiated. In summary, both monkeys completed about 840 trials per session (Monkey F: 843 ± 48; Monkey G: 841± 37)

### Neurophysiology

Subjects were implanted with recording chambers at the front left, targeting posterior dACC (Brodmann area 24c, subject F: 23 mm anterior to the interaural plane, 13 mm medial to the sagittal midline, 0 degree tilted; subject G: 27.8 mm anterior, 14.6 mm medial, 40 degrees tilted), which has been reported to respond to more cognitive aspects of learning compared to the anterior part^36^. Single unit activities were recorded using linear multichannel probes (Poly 2 or edge probe, Neuronexus, Ann Arbor, Michigan) with 32 channels (spanning two lanes (16 each) per line) with average spacing of 100*μm* between every adjacent channel (1*mΩ*). To determine the recording location, we performed structural imaging using a 3T MRI scanner (Siemens, Munich, Germany) prior to the chamber implant. The resulting scan of the brain structure allowed us to plan the implant trajectory and stereotaxic coordinates from the Brainsight (Rogue Research, Montreal, Quebec, Canada). After the surgery, we performed an additional structural MRI scan to verify the location of the chamber as well as match each location in the chamber with the targeting part of the brain. Every session, a recording probe was penetrated into the brain by the motorized micro drive system (NaN Instruments, Nazareth, Israel, recording range: anterior to AP monkey F: 22.8-26.9mm, monkey G: 22.9-29.2mm). Depth of penetration to dACC was determined based on MRI, further verified by empirical observation of high frequency spikes first, then silence and lastly medium frequency spikes when the probe went through the cortical area (FEF or premotor cortex), then white matter and dACC. Raw spikes were amplified and digitized (RHD 32 channels processor, Intan Technologies, Los Angeles, California), high-pass filtered (300-6000 Hz) and then sampled at 30 kHz by the Open Ephys data acquisition system (version 2.2, Cambridge, Massachusetts), which synchronized the data stream from brain, licking and all behavior variables. Both the raw LFP and filtered neural signals were saved in binary data format.

We used Kilosort (version 2.5^37^ and version 4^38^) for spike sorting. The procedure is as follows: filtered spike binary files were imported to both versions of Kilosort in parallel. Each cluster was then manually inspected, merged or split based on overall spikes count, waveform, inter-spike interval (ISI) histogram and spikes feature projected in the low dimensional space. Then, we plotted a raster map for each cluster and did final examination based on if firing rates and spike amplitudes drifted during the whole session. Only the units that passed all criterions were included. Lastly, we compared sorting performance between two Kilosort versions and picked the one that yielded more neurons. Overall, we collected 1072 units from dACC (F: 426; G:636).

### Behavioral analysis

To quantify how strongly subjects ordered stimuli under two environments, we implemented the procedure as before^21^. Briefly, we assumed preferences towards each stimulus were distributed along a vectoral continuum, parametrized by its *z*-score *μ* and uncertainty *σ*. The probability of choosing *X* when paired with *Y* was given by:

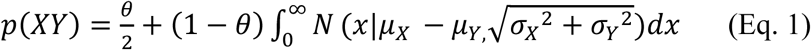

This shows that chosen probability depends on the difference and overall uncertainty between *z*-scores of both stimuli, as well as the degree of being insensitive to the preferences by randomly responding (θ). We constructed the model within the Bayesian updating framework and fitted the model to the data only from the testing phase. Best-fit estimates of *μ* and *σ* were obtained using Markov chain Monte Carlo (MCMC) and implemented in Stan^29^. Afterwards, we performed linear regression over all *z*-scores sorted in an ascending order. This can let us use the slope as a metric to reflect the ordering preference (OP) and denote the subjective rank E-A for each pictorial set, particularly the U set. Meanwhile we can contrast the OP for L versus U to calculate OP_diff_ to examine the strength of behavioral differentiation between the two sets.

We also simulated data of random responses 80 times for each subject to acquire the “baseline” slopes, which served as the null model against which the actual slopes were compared. This provided a more conservative null hypothesis, as even random responding is expected to favor some items over others by pure chance during any given trial, resulting in slightly positive baseline slope.

### Statistical analysis on neural data

#### Neural activity visualization

To directly see the neural activity, we constructed the PSTH for each neuron with the activity aligned on the time point of stimulus presentation, smoothed by a gaussian kernel (kernel width: 240ms) and then normalized by the baseline activity defined by the averaged firing rates within the 500ms before the stimulus display.

#### Time resolved Poisson General linear model (Poisson GLM)

To look at learnability modulation, we constructed the linear mixed-effect model as depicted:

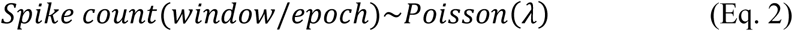

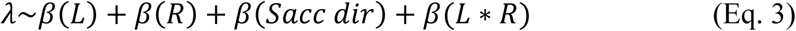

We first extracted the spikes from the testing phase only since behavior was more stable compared to training (**Fig 1G, H, S1**). Then we prepared the spike data in two ways. First way is to count the spikes from four major epochs of the trial: 800ms before stimulus on; stimulus on to choice; choice to feedback and 2000ms after the feedback. Second way is to better visualize the dynamics of encoding. We slid a time window (window size=300ms) from 800ms before stimulus on until 3700ms after with 50ms per stride. We regressed the spike counts within each epoch/window on the learnability(L), reward outcome(R) their interaction and saccade direction, using the `sm.GLM(family=sm.families.Poisson())` function from the statsmodels python module. Since in the learnable set, choosing higher rank always incurred higher reward, we did not include rank to prevent collinearity among predictors. Besides reporting the *β* and *p*-value from each regressor, we also calculated the coefficient of partial determination (CPD)^39^, which computed for *i*^th^ predictor in the *k*^th^ time window:

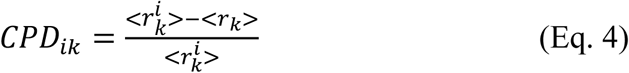

where *r*_*k*_ denotes the squared norm of the residual vector and *r*^*i*^ denotes the same over the reduced model when regressor *i* is removed. In other words, CPD measures the normalized contribution of the variance of spike counts across trials by each regressor. We also performed the random permutation test 50 times by shuffling each regressor, running Poisson GLM and calculating CPD independently. We compared the CPDs from the true and shuffled models using a non-paired t test to verify if the CPD is significant. Before regression, we constructed a confusion map over all regressors to ensure neither two regressors were highly correlated. Indeed, there were no two predictors who showed correlation strength higher than 0.7 in more than 10% of the sessions.

We also implemented similar Poisson GLM within each list to examine effects by outcome:

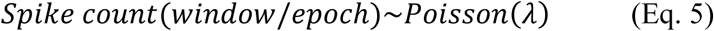

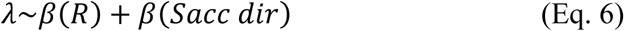

All the other procedures followed the previous regression.

### Decoding analysis

To investigate how the population as a whole represented learnability and outcome, we first select sessions with more than 10 units simultaneously recorded. Then we counted the spikes from four major epochs of the trial similarly to the epoch-wise GLM. For each epoch, we trained a logistic classifier on the trial types and outcome separately from the population activities within each monkey as well as all data pooled, using Matlab built-in function `fitclinear` (Logistic learner, L2 regularization). We evaluated the decoding performance based on five-fold cross validation accuracy. Performance was compared with averaged accuracies from the control group where labels were shuffled across trials 100 times.

## Acknowledgments

We thank Dr. Fabian Munoz and Yvoone Li for technical help with electrophysiology. We also thank Dr. Liam Paninski and Dr. Stuart Firestein for valuable discussions on analysis and interpretation the project.

## Funding

This work was supported by grant NIH-R01MH111703 from the National Institutes of Health (VPF) and NIH-R34NS137420 (VPF and JG)

## Data sharing plan

Code and data are available from the first author upon request.

## Author contributions

The study was designed and the manuscript was prepared by YJ, GJ, JG & VPF. Data were collected and analyzed by YJ. The authors declare no competing interest.

## Notes

### Competing Interest Statement

The authors have declared no competing interest.

